# Polygenic transcriptome risk scores improve portability of polygenic risk scores across ancestries

**DOI:** 10.1101/2020.11.12.373647

**Authors:** Yanyu Liang, Milton Pividori, Ani Manichaikul, Abraham A. Palmer, Nancy J. Cox, Heather Wheeler, Hae Kyung Im

## Abstract

Polygenic risk scores (PRS) are on course to translate the results of genome-wide association studies (GWAS) into clinical practice. To date, most GWAS have been based on individuals of European-ancestry, meaning that the utility of PRS for non-European populations is limited because SNP effects and LD patterns may not be conserved across populations. We hypothesized that cross population prediction at the level of genes rather than SNPs would be more effective, since the effect of genes on traits is likely to be more highly conserved. Therefore, we developed a framework to convert effect sizes at SNPs into effect sizes for genetically predicted transcript abundance, which we used for prediction in non-European populations. We compared this approach, which we call polygenic transcriptome risk scores (PTRS), to PRS, using data from 17 quantitative traits that were measured in multiple ancestries (European, African, East Asian, and South Asian) by UK Biobank. On average, PTRS using whole blood predicted transcriptome had lower absolute prediction accuracy than PRS, as we expected since not all regulatory processes were captured by a single tissue. However, as hypothesized, we found that in the African target set, the portability (prediction accuracy relative to the European reference set) was significantly higher for PTRS than PRS (p=0.03) with additional gain when transcriptomic prediction models ancestry matched the target population (p=0.021). Taken together, our results suggest that using PTRS can improve prediction in underrepresented populations and that increasing the diversity of transcriptomic data may be an effective way to improve portability of GWAS results between populations and help reduce health disparities.

## Introduction

Polygenic risk scores (PRS) for a variety of traits are increasingly becoming accurate enough to be useful for clinical practice, realizing the longstanding goal of personalized medicine. PRS for coronary artery disease (CAD) have been shown to provide prediction that has been compared to monogenic mutations of hypercholesterolemia (Khera et al., 2018). In practice, PRS may impact a larger proportion of patients compared to monogenic mutations; for example, PRS for CAD provide potentially actionable information for 8% of the population (for whom the risk increases by three-fold) whereas known monogenic mutations are only informative for about 0.4% of patients. However, a major limitation of this approach is that PRS developed in one human ancestry group do not perform well in other ancestry groups, limiting their utility and exacerbating already severe health disparities (Curtis, 2018; Martin et al., 2019). This problem is being addressed by large efforts such as Human Heritity and Health in Africa (H3Africa) Choudhury et al. (2020), Million Veterans Project (Gaziano et al., 2016), AllofUs (of Us Research Program Investigators, 2019) and TOPMED (Taliun et al., 2019) that are recruiting individuals from diverse ancestry groups.

However, these efforts are time consuming, enormously expensive and will have to be repeated at scale, for numerous traits, across numerous ancestry groups. Therefore, methods that can use GWAS results from one population for prediction in other ancestry groups is highly desirable. Analysis of GWAS conducted in different populations suggested that a considerable fraction of causal SNPs are shared across populations (Shi et al., 2020). This suggests that efforts to develop methods that transfers knowledge across populations can provide a cost effective ways to improve prediction in underrepresented ancestry groups. Many loci identified by GWAS are thought to exert their effects by regulating gene expression. Motivated by this mechanistic insight, multiple eQTL studies have been performed over the last decade (The GTEx Consortium, 2020; Võsa et al., 2018). The GTEx consortium, for example, has sequenced mRNAs samples from 50 tissues across the human body from more than 900 donor individuals. PrediXcan (Gamazon et al., 2015) and other TWAS methods (Gusev et al., 2016; Hu et al., 2019) leverage these reference transcriptome datasets to train prediction models of gene expression levels and correlate the genetically predicted gene expression levels with complex traits to identify causal genes. Given the common biology of human disease across populations and the mediating role of the transcriptome, we hypothesized that that prediction at the level of estimated transcript abundance rather than SNPs might be more effective across populations.

Therefore, we propose the polygenic transcriptomic risk score (PTRS) as a gene-based complement to the PRS that has the potential for superior portability across human ancestry groups. One advantage of PTRS is that the smaller number of features (tens of thousands of genes rather than millions of SNPs), means that optimizing the parameters to build PTRS is more manageable than PRS. Another advantage of PTRS is that training transcriptome prediction models requires much smaller samples than training PRS, and can then be used for prediction of many different traits. Furthermore, training data for non-European individuals are becoming increasingly available. Finally, because PTRS is gene-based, it is inherently more biologically interpretable than PRS.

In this paper, we explored the advantages of PTRS using the UK Biobank (UKB), which provides genotype and phenotype data in half a million individuals (Bycroft et al., 2018). Although the majority of participants in UKB are of European-descent, several thousand individuals of non-European descent are also available, and could be used to compare prediction by PRS and PTRS across ancestries. We started by testing whether the genetically predicted transcriptome could explain a sizeable portion of trait heritability and whether matching the transcriptome training and risk score testing populations’ ancestry would be beneficial. Then, we built PRS and PTRS for a range of complex traits and compared their prediction accuracy and portability across populations.

## Results

Before describing the results we define and clarify some terminology. In this paper, there are two types of prediction: 1) gene expression level prediction from genotype data and 2) complex trait prediction using PRS or PTRS. PRS uses genotype data directly and PTRS uses linear combinations of genotypes representing predicted gene expression levels. To simplify exposition, we will only use the term *training* for the calculation of weights for predicting gene expression levels using genotype data. The *training* of transcriptome (gene expression levels) prediction weights had been performed previously and we simply downloaded them from predictdb.org. When we estimate optimal weights for PRS and PTRS, we will use the terms *building* or *constructing*. We performed the *building* of PRS and PTRS using the *discovery* set. The testing of the risk scores, PRS and PTRS, were performed in what we call the *target* sets. For the remainder of the paper, we will refer to individuals by their ancestry and drop the - descent suffix. Unless otherwise clarified, we will use the term transcriptome to mean the set of predicted expression levels of genes. GTEx EUR transcriptome should be interpreted as the set of predicted gene expression levels using weights trained in European samples from GTEx. Similarly, MESA EUR transcriptome, will refer to the predicted transcriptome using weights trained with the MESA European samples. MESA AFHI transcriptome will refer to the predicted transcriptome using weights trained with a combination of African American and Hispanic individuals from the MESA study.

### Experimental setup

An overview of the experimental setup describing the discovery, training, and target sets used in the paper is shown in Fig.1. We randomly selected 356,476 unrelated Europeans in the UK Biobank for the discovery set. For testing the performance of risk scores, we constructed 5 target sets with participants of various ancestries in the UK Biobank. We used 2,835 African, 1,326 East Asian, and 4,789 South Asian individuals for the non European target sets. We also reserved two randomly selected sets of 5,000 Europeans as additional target sets. One was selected as the EUR reference set and the second European target was used as a test set to assess the the variability of the results within the same ancestry.

**Figure 1:**
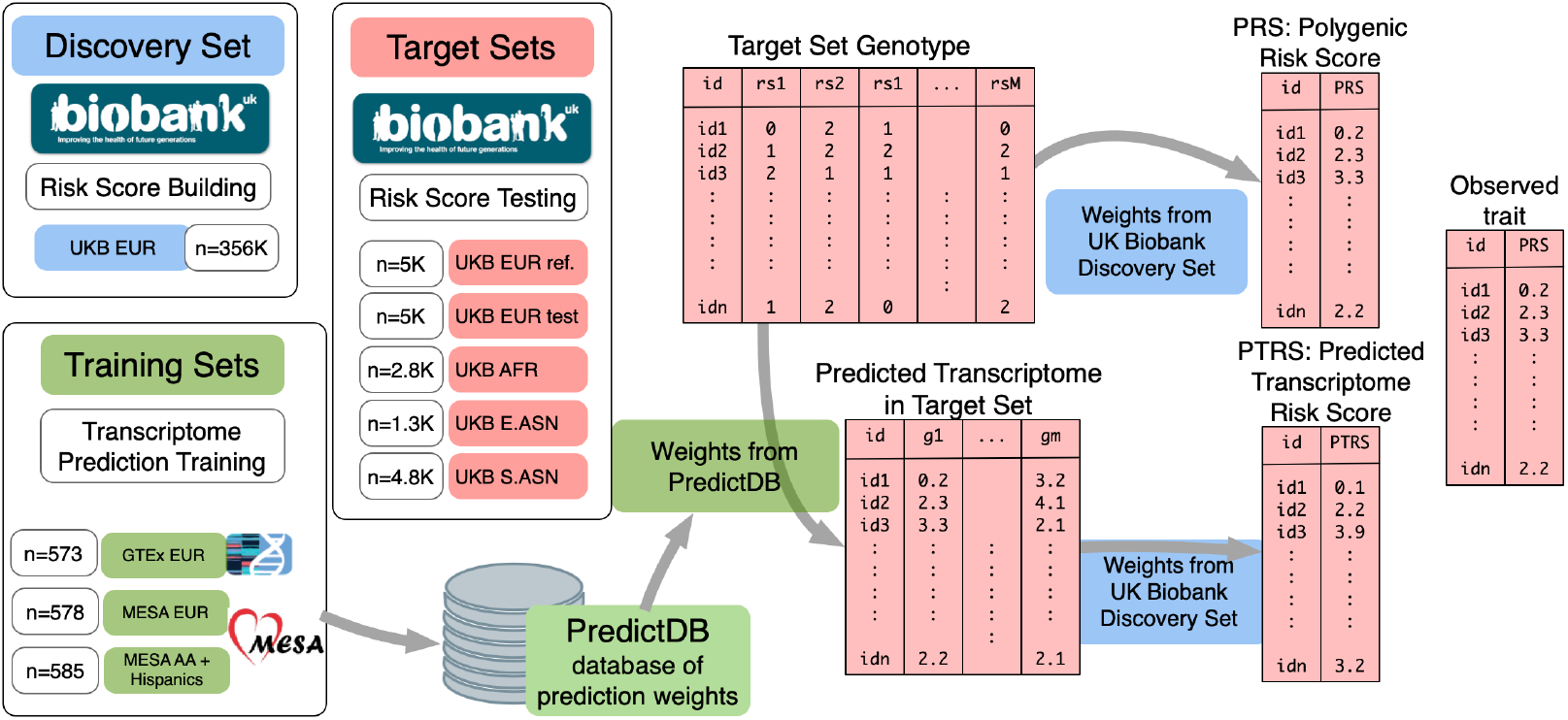
Experiment setup. This figure summarizes the experimental set up used for testing the portability of PRS and PTRS across populations. The weights for calculating PRS and PTRS were estimated in the *discovery set*, which consisted of 345K randomly sampled individuals of European-descent from the UK Biobank. The *training* sets where the weights for the prediction of transcriptomes were computed are shown in green. We downloaded the weights trained previously from predictdb.org. We sampled 5 *target sets* from the UK Biobank for testing the risk scores: two randomly sampled sets of European-, one African-, one East Asian-, and one South Asian-descent individuals. For each of the 5 *target* sets, predicted transcriptomes were calculated using the weights trained in each of the three *training* sets: GTEx EUR, MESA-EUR, MESA-AFHI.

For predicting the transcriptome, we downloaded prediction weights from multiple ancestries collected in predictdb.org. The first set of models had been trained in European individuals from the GTEx v8 release (Barbeira et al., 2020a) in whole blood and 9 other tissues chosen by had been large sample size. The second set of models had been trained using monocyte samples of Europeans, African Americans, and Hispanics from the MESA cohort (Mogil et al., 2018).

For our tests, we focused on the 17 anthropomorphic and blood phenotypes used by Martin et al. (Martin et al., 2019).

### Predicted transcriptome captures a significant portion of trait variability

To assess the feasibility and to quantify the potential for PTRS to predict human traits, we calculated the proportion of variance explained by the predicted transcriptome assuming random effects of gene expression levels. The approach is analogous to standard SNP-heritability estimation (Yang et al., 2010) where the “predicted expression relatedness matrix” is used instead of the genetic relatedness matrix. In this section, we calculated the predicted transcriptome using the GTEx EUR weights using the European target set genotype data. Using these predicted expression levels, we calculated the “predicted expression relatedness matrix” (instead of the genetic relatedness matrix) and applied the standard restricted maximum likelihood estimation to calculate the proportion of variance explained by the predicted transcriptome.

Since the PVE varies depending on the underlying heritability of the trait, we also calculated the proportion of SNP heritability explained by the predicted transcriptome as the ratio of PVE and heritability. We term this values “regulability” (Barbeira et al., 2020a). For the SNP-heritability we used publicly available heritability estimates based on LDSC regression (Bulik-Sullivan et al., 2015) in UK Biobank Europeans for the same set of phenotypes (see details in section 1.7).

As shown in Fig.2A, in the European target set, GTEx EUR whole blood based predicted transcriptome captured on average 20.6% (s.e.=2.1%) of the trait variability. This result is largely consistent to the estimates reported previously using a different approach (Yao et al., 2020).

**Figure 2:**
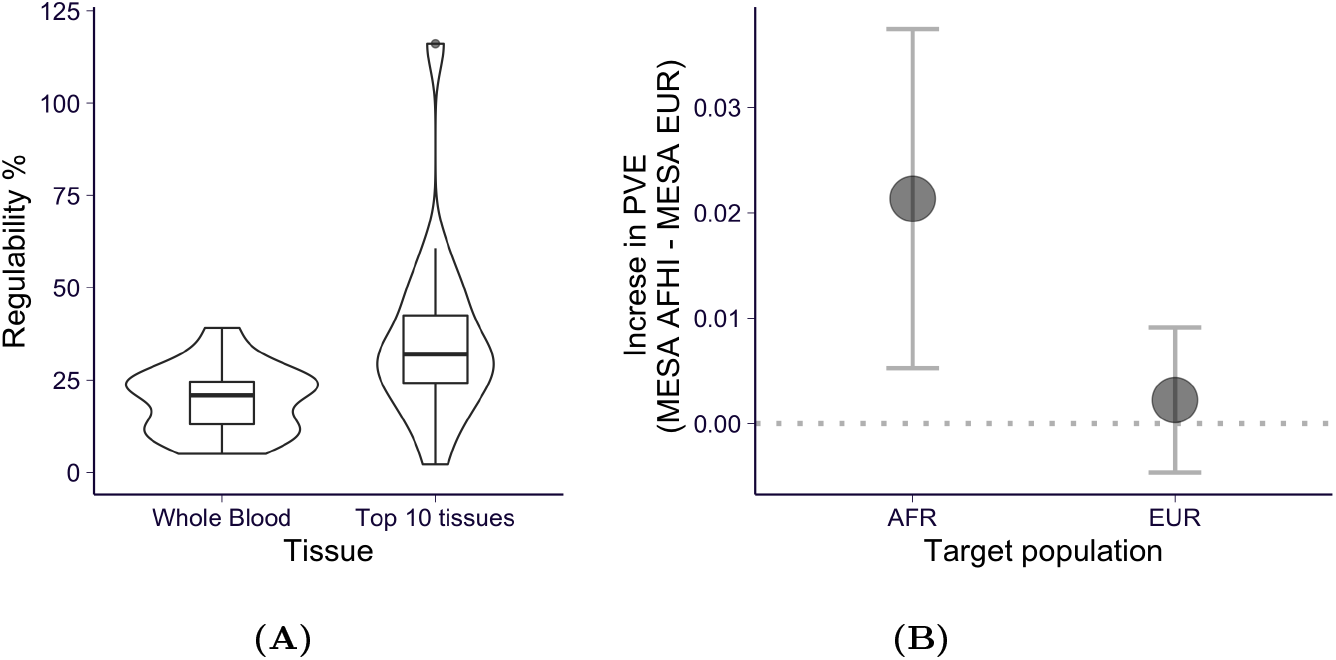
Proportion of variance explained (PVE) by the predicted transcriptome. **(A)** shows the ratio of PVE (the proportion of phenotypic variation explained by the predicted transcriptome) of GTEx EUR transcriptome model over the chip heritability using whole blood on the left and using 10 tissues on the right. **(B)** shows the mean of the difference between the PVE by the predicted MESA AFHI transcriptome and the PVE by the MESA EUR transcriptome. For the African set, the MESA AFHI transcriptome explains more variance that the MESA EUR transcriptome. In the European set, the difference between the two transcriptomes is not significant. The vertical bars show the 95% confidence intervals estimated with paired t-test.

### Aggregating predicted transcriptomes in multiple tissues increases the PVE

To explore ways to increase the proportion of variance explained (PVE), we calculated the proportion explained collectively by the transcriptome predicted in 10 tissues, including muscle, adipose, tibial artery, breast, lung, fibroblast, lung, tibial nerve, and skin, with sample sizes ranging from 337 to 602 (Supplementary Table S3). As anticipated, we found that, collectively, the predicted transcriptomes in 10 tissues explained a larger portion of heritability: on average 34.4% (s.e. = 3.3%) of the heritability corresponding to a 48% increase relative to whole blood alone. This result suggests that adding transcriptomes from multiple tissues will improve predictions in general.

### Matching training and target ancestries increases the proportion of variance explained

Next, we examined whether using transcriptomes trained in a population genetically closer to the target set could explain a larger proportion of the trait variance. For this, we took advantage of the availability of trans-ancestry transcriptome prediction models from the MESA cohort (Mogil et al., 2018). One of them (MESA-EUR) was trained in a European population and the other one (MESA AFHI) was trained in a combination of African American and Hispanic populations (Supplementary Table S3). We decided to use the combined (African American and Hispanic) transcriptome prediction since the similarity of the sample sizes (578 vs 585) would make the comparison with the European trained models more fair.

We found that (Fig.2B) in the African target set, using the ancestry matched MESA AFHI transcriptome yielded a significantly higher (2.1% with s.e. = 0.8%) proportion of variance explained than when using the MESA EUR transcriptome. For the European target set, the difference between using the MESA AFHI or the EUR transcriptomes was not significantly different from 0.

### Building PRS and PTRS

After having determined that it is possible to capture a significant portion of trait variability using predicted transcriptome and that matching the training and target set ancestries can increase the portion explained, we proceeded to build the PRS and PTRS in our discovery set (356K Europeans from the UK biobank).

We built PTRS weights using elastic net, a regularized linear regression approach, which selects a sparse set of predicted expression features to make up the PTRS. For PRS weights, we used the standard LD clumping and p-value thresholding approach (see details Methods section section 1.3).

We quantified the prediction accuracy in each target set using the partial *R*^2^ 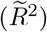, which provides a measure of correlation between predicted and observed outcomes with the added benefit of taking covariates into account (see details in section 1.10).

All the weights were calculated in the discovery set, however, to boost the prediction performance across the board, we performed an additional tuning step in the target populations. This was done for all scores (PRS and PTRS) in each target set so that the comparison remains fair. For PRS, we chose the p-value threshold that yielded the highest 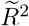 in each target set. For PTRS, we pre-computed weights for a range of regularization parameters in the discovery set and chose the parameter that maximized the 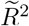 in each target set.

### PTRS prediction accuracy outperforms expectation given their PVE explained

Tested in the European target sets with European training set transcriptome models, the mean prediction accuracy of PTRS (GTEx EUR based) was lower than the accuracy of PRS (paired t-test p = 0.03) as shown in Fig.3A for the 17 traits. However, PTRS performance was much higher that what could have been expected given that predicted expression only explained about a fifth of the trait variation explained by typed and imputed SNPs. This better than expected performance indicates that integrating predicted transcriptomes and other omics is a promising avenue to improve PRS performance in general.

**Figure 3:**
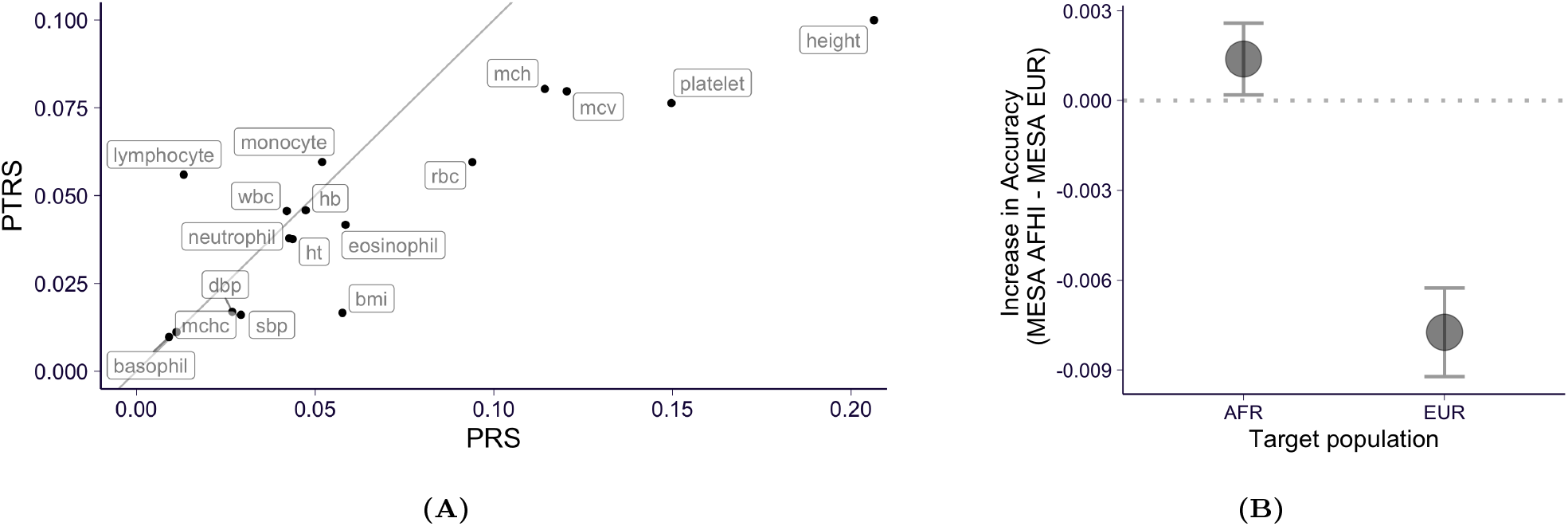
Prediction accuracy of predicted transcriptome risk scores (PTRS). **(A)** Prediction accuracy, measured by partial 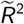, of PTRS (on y-axis) compared to the accuracy of PRS (on x-axis). Given the fact that the PVE by predicted expression was on average 20.1% of the heritability, PTRS is performing much better than expected. **(B)** The difference in the prediction accuracies bewteen MESA AFHI and MESA EUR models for the set of African samples and European samples. Matching training and target set ancestries leads to improved prediction accuracy: AFHI transcriptome yields better prediction accuracy in the African target set and EUR transcriptome yield better prediction accuracy in the European target set.

### Matching training and target ancestries improves prediction accuracy

To test whether matching the training and target ancestries would improve the PTRS prediction accuracy, we examined the difference between using the African transcriptome (MESA AFHI) vs the European transcriptome (MESA EUR). As hypothesized, the PTRS based on the MESA AFHI transcriptome yielded accuracy higher than the PTRS based on the MESA EUR transcriptome when the target set was African. Similarly, for the European target set, the European transcriptome based PTRS had better accuracy than the AFHI transcriptome based ones. Fig.3B shows the small but significant difference (*R*^2^(AFHI) -*R*^2^(EUR)), which was positive (0.14%, s.e.= 0.06%) for the African target set and negative for the European target sets (−0.77% s.e.=0.08%), consistent with the positive effect of matching the ancestries.

To avoid differences due to having different number of predicted genes, PTRS were built using only the genes that were present in both training sets, EUR and AFHI. See details in Methods section section 1.8.

### PTRS improves portability into the African population

To test our hypothesis that PTRS can generalize more robustly across populations than the standard PRS, we defined ‘*portability*’ as the predictive accuracy in each population relative to the European reference target set (EUR ref.). This is calculated as the ratio of the 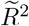 in the target population divided by the 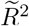 in the European reference target set. Thus, by definition the portability in the European reference set is 1.

Consistent with reports by (Martin et al., 2019), the portability of PRS degrades with the genetic distance to the European discovery set as shown in gray in Fig.4A. The portability of PTRS (shown in orange) also decreases with genetic distance to the discovery set, with the African target sets showing the largest loss of accuracy, as expected. However, we also observed that the portability of PTRS in the African target set was significantly higher than the portability of PRS (paired t-test p=0.03). These results provide strong proof of principle that integrating predicted transcriptome as done with PTRS has the potential to improve portability of risk scores across populations.

**Figure 4:**
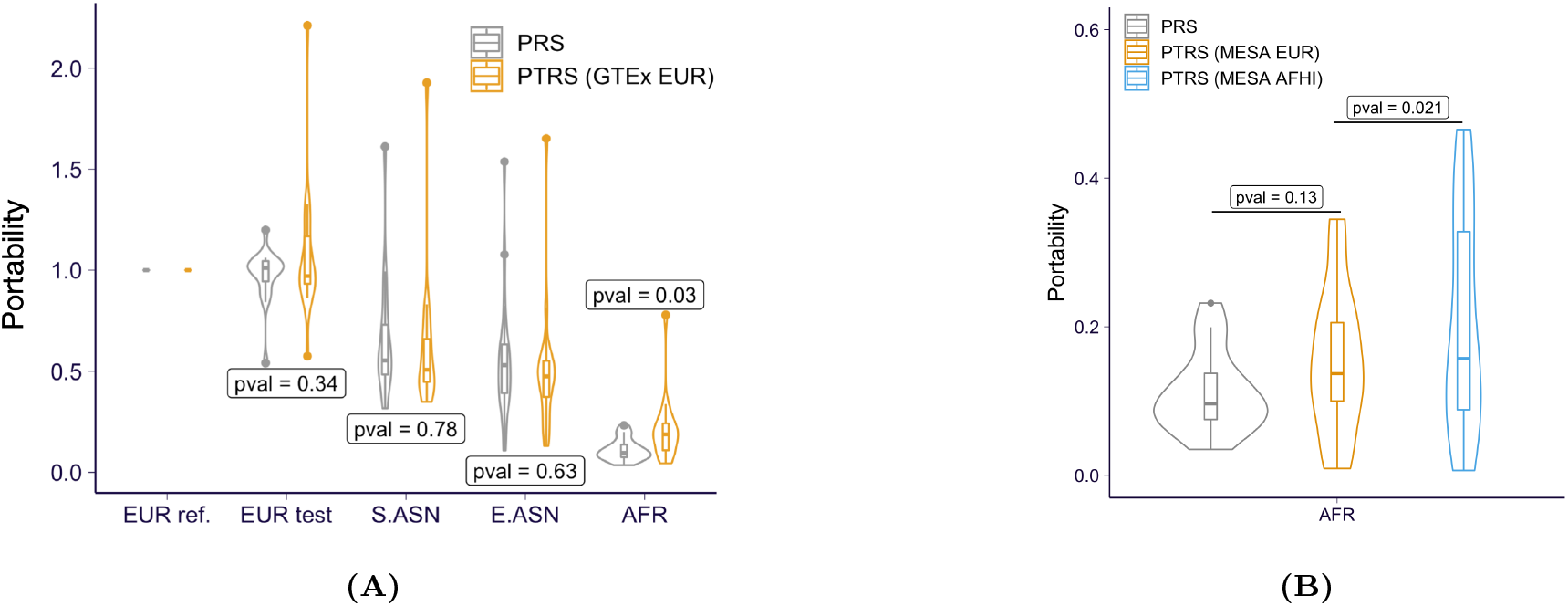
Portability of PTRS for 17 quantitative phenotyes in UK Biobank. **(A)** The portability of PTRS trained and calculated using GTEx EUR whole blood samples are shown in yellow with the PRS shown in gray (left panel). ‘EUR ref.’ set is used as the reference population in the calculation of portability (section 1.11) so that the portability is always 1. **(B)** The right panel shows the portability in the AFR target set using the MESA transcriptome models trained in EUR and AFHI sets. We observed the trend that PTRS has better portability using EUR transcriptomes both from GTEx and MESA. The gain in the AFR set is even higher when AFR transcriptomes are used for the PTRS.

In the European test set, we observed quite a bit of variability in the portability, ranging from 0.54 to 2.21, despite the fact that both European target sets were randomly sampled from the same European UK Biobank participant set. As expected, the median portability in the second EUR target set is centered around 1.

Finally, we used the MESA EUR and AFHI models to assess the potential improvement in portability when matching training and target set ancestries. As shown in Fig.4B, PTRS based on AFHI transcriptome has significantly higher portability than the MESA-EUR transcriptome-based PTRS (paired t-test p=0.021). As an additional evidence of improved portability of PTRS in general, we replicated the higher portability of the EUR-based PTRS compared to PRS using an independent training set (MESA vs GTEx as described above).

Taken together, our results provide support to our hypothesis that PTRS can transfer more robustly than PRS, which can be improved by using ancestry matching transcriptomes. Also they suggests that adding transcriptomes predicted in other tissues and other omics data can further improve PRS generalizability.

## Discussion

In this paper, we showed that informing genetic risk score building using genetically determined gene expression traits as intermediate predictors as implemented with PTRS can lead to predictors that are more portable across populations, especially if matched ancestry transcriptomes are used.

We found that the total trait variation that can be explained via predicted transcriptomes range from 20.6% (using whole blood) to 34.4% (with a broader sets of tissues) of the SNP heritability, i.e. the total variation that can be explained using common SNPs. Promisingly, the actual predictors built on predicted transcriptomes had performances that were much higher than the expected 20.6% of the PRS performance. The predicted transcriptome tended to be more predictive if it was trained with individuals from the same ancestry stressing the advantages of collecting transcriptome reference data in diverse populations.

We found that the portability of PTRS was significantly higher than the portability of PRS in the African target set, the most affected by the Eurocentric bias in GWAS studies, with further gains when the transcriptome was trained in matched ancestry samples.

Our results show that investing in multi-omic studies of diverse populations may be a cost effective way to reduce current genomic disparities by taking better advantage of existing GWAS studies. Developments of methods to optimally combine PRS and PTRS should be encouraged.

Our study points to promising strategies to improve risk prediction in general but it also has several limitations. First, PTRS are based on prediction models of gene expression traits which we estimated to account for less than a third of the total variability in the complex traits considered here. We expect this limitation to be mitigated as additional transcriptome reference sets as well as other omics data covering mediating mechanisms missed in current models. Second, we used single tissue prediction models for most of the analysis in this paper, which captured a fifth of the variation in the complex traits here. We will develop approaches to integrate multiple tissue models. Third, weights for PRS were calculated using GWAS summary results (thresholding and pruning method) whereas PTRS weights were calculated using individual level data due to computational considerations. Future analysis will be performed using individual level data for PRS by using biobank-scale ready elastic net approaches such as (Qian et al., 2020). Fourth, higher quality prediction models of the transcriptome in non-European ancestries are limited. Here we used predictors trained in monocyte samples assayed with older array technology. Multiple ancestry models are currently being generated by us and other groups. For example, the MESA TOPMED project has assayed RNAseq, protein, methylation, and metabolomics data in African Americans, Hispanics, and Asian ancestry individuals which will allow the development of improved prediction models.

## 1 Material and Methods

### 1.1 Obtaining individuals and phenotypes from UK Biobank

We used data from UK Biobank data downloaded on July 19 2017. We excluded related individuals and the ones with high missing rate or other sequencing quality issues. As covariates, we extracted age at recruitment (Data-Field 21022), sex (Data-Field 31), and the first 20 genetic PCs. The ancestry information of individuals was obtained from Data-Field 21000 and we kept individuals labelled as ‘British’, ‘Indian’, ‘Chinese’, or ‘African’ (according to Data-Coding 1001: http://biobank.ctsu.ox.ac.uk/crystal/coding.cgi?id= 1001). Throughout the paper, we labeled ‘British’ individuals as EUR, ‘Indian’ individuals as S.ASN, ‘Chinese’ individuals as E.ASN, and ‘African’ individuals as AFR. The measurements of the 17 quantitative phenotypes (as shown in Supplementary Table S1) across all available instances and arrays were retrieved. The data retrieval described above was performed using ukbREST (Pividori and Im, 2019){pividori:2019} with the query YAML file available at https://github.com/liangyy/ptrs-ukb/blob/master/output/query_phenotypes.yaml.

If one individual has multiple measurements for the same phenotype (in more than one instances and/or more than one arrays), we collapsed multiple arrays by taking the average and we aggregated measurements across multiple instances by taking the first non-missing value. Individuals with missing phenotype in any of the 17 quantitative phenotypes or covariates were excluded.

### 1.2 Quality control on self-reported ancestry

To ensure the quality of ancestry label, we removed individuals who deviate substantially from the population that they were assigned to. Specifically, for population *k* among the 4 populations (EUR, S.ASN, E.ASN, and AFR), we treated the distribution of the individuals, in the space of the first 10 PCs, as multivariate normal. And we calculated the observed population mean 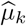 and covariance 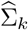 accordingly. Then, for each individual *i* in population *k*, we evaluated the “similarity” *S_ik_* to the population *k* as 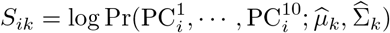 Intuitively, if an individual has genetic background differing from is the assigned population, the corresponding *S_ik_* will be much larger than others. So, we filtered out individuals with *S_ik_* ≤ −50 in the assigned population *k*. This cutoff was picked such that 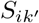 for any un-assigned population 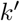 has 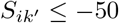 for all individuals.

The number of individuals remained after data retrieval and ancestry quality control is listed Supplementary Table S2.

### 1.3 Performing GWAS and building LD-clumping based PRS models

We built PRS using the genotypes and phenotypes of the individuals in the discovery data set (the details of data splitting is described in section 1.11). We performed GWAS (linear regression) using linear_regression_rows in hail v0.2 where we included covariates: first 20 genetic PCs, age, sex, age2, sex × age, and sex × age2. In the GWAS run, we excluded variants with minor allele frequency *<* 0.001 and variants that significantly deviate from Hardy-Weinberg equilibrium (p-value *<* 10^*−*10^). And the phenotype in their original scales were used.

To obtain relatively independent associations for PRS construction, we run LD clumping using plink1.9 with option `–clump-clump-p1 1 -clump-r2 0.1 –clump-kb 250`. This command extracted genetic variants in the order of their GWAS significances and excluded all variants having *R*^2^ *>* 0.1 to or 250 kb within any variants that have already been included. The PRS was constructed on the basis of the set of variants obtained from the LD clumping along with the marginal effect size estimated in GWAS run. Specifically, we calculated PRSs at a series of GWAS p-value thresholds: 5 × 10^*−*8^, 10^*−*7^, 10^*−*6^, 10^*−*5^, 10^*−*4^, 10^*−*3^, 0.01, 0.05, 0.1, 0.5, and 1. In other word, at threshold *t*, the PRS for individual *i* was calculated as

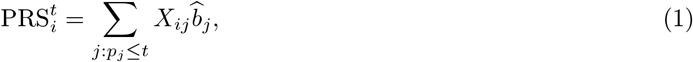

where *X_ij_* is the effect allele dosage of variant *j* in individual *i* and 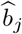 is the estimated effect size of variant *j* from GWAS run.

At the testing stage, given the genotype of an individual, we calculated the PRS of the individual using eq. (1).

### 1.4 Computing the predicted transcriptome

We computed predicted gene expression for all individuals passing filtering s teps a nd q uality c ontrol. We utilized two sets of prediction models: 1) CTIMP models (proposed in (Hu et al., 2019)) trained on GTEx v8 EUR individuals (Barbeira et al., 2020b){barbeira:2020}; and 2) elastic net models which were trained on Europeans (EUR) or African Americans in combination with Hispanics (AFHI) (Mogil et al., 2018){mogil:2018}. The sample size and tissue informations of the prediction models are listed in Supplementary Table S3.

### 1.5 Estimating PVE by predicted transcriptome of a single tissue

To get a sense on the predictive power of predicted transcriptome on the phenotypes of interest, we estimated the proportion of phenotypic variation that could be explained by the predicted transcriptome in aggregate.

Specifically, we assume the following mixed effect model (for individual *i*).

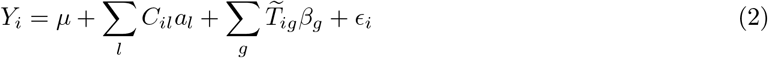

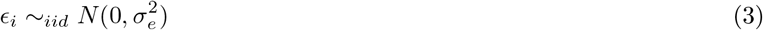

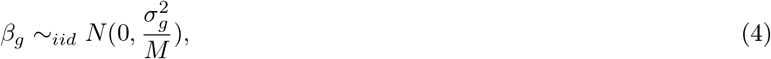

where *M* denotes the number of genes, *C_il_* is the *l*th covariate,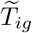 is the inverse normalized predicted expression for gene *g*, and *Y_i_* is the observed phenotype. By inverse normalization, we converted the predicted expression 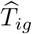 to 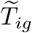 by 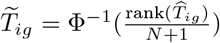 within each gene *g* where *N* is the number of individuals and ‘rank’ is in increasing order. So that we have 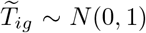. The parameters of the model were estimated using hail v0.2 `stats.LinearMixedModel.from_kinship` with *K* matrix being set as 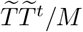. And PVE is calculated as 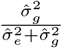 The same set of covariates as section 1.3 were used.

The PVE estimation was performed for each transcriptome model and population pairs. For non-European populations, all individuals were included in the analysis. We randomly selected 5,000 EUR individuals for the analysis.

### 1.6 Estimating PVE by predicted transcriptome of multiple tissues

The genetic effects on the complex trait can be mediated through the regulation of expression in different tissues so that including predicted transcriptomes in multiple tissues could potentially improve the prediction performance. To test this idea, we performed PVE analysis as described in section 1.5 using predicted expression in 10 GTEx tissues (listed in Supplementary Table S3). To account for colinearity issues induced by the high correlation of predicted expression among tissues, we preselected linearly independent ‘eigen-predicted expression’ traits using singular vectors of the predicted expression data. This approach is similar to the one used for combining PrediXcan association in multiple tissues (Barbeira et al., 2019). The PVE was calculated using a mixed effects model similar to eq. (2) where the expression traits were replaced by the eigen-predicted expression traits.

Briefly, the eigen-predicted expression traits were calculated as follows. For each gene *g*, let 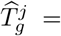 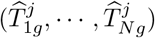 denote the predicted expression (with standardization) of *g* in tissue *j* across individual *i* = 1, · · ·, *N*. By collecting 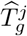 for all tissues which have prediction model for gene *g* (suppose there are *J* of them), we have a matrix 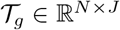 where columns correspond to tissues. To remove the colinearity in the columns of 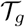 by keeping linearly independent predictors, we used the first *K* left singular vectors of 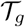 with *K* selected as follows. We performed PCA on 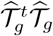 and any *k*th PC 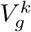 was excluded if λ_k_/λ_max_ ≤ 1/30. The leading *K* left singular vectors (up to a scaling factor) of 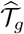 q was reconstructed as follow.

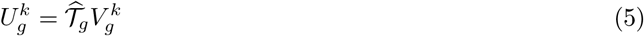

We further inverse normalized 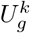 (as described in section 1.5) resulting in 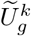, which were plugged into

### 1.7 Retrieving publicly available heritability estimates

We retrieved chip heritability estimates by LDSC regression (Bulik-Sullivan et al., 2015) from https://nealelab.github.io/UKBB_ldsc/downloads.html where we downloaded the file https://www.dropbox.com/s/8vca84rsslgbsua/ukb31063_h2_topline.02Oct2019.tsv.gz?dl=1. These estimates were calculated with GWAS summary statistics obtained from UK Biobank Europeans with inverse normalized phenotypes. We extracted phenotypes of interest by their UKB Field ID. And we used columns h2_observed and h2_observed_se as the estimated value and standard error of the heritability estimation.

### 1.8 Building PTRS models using elastic net

For each of the 17 quantitative phenotypes, we trained elastic net model to predict the phenotype of interest using the predicted transcriptome (in a single tissue) as features. The same set of covariates as described in section 1.3 were used. Let 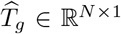 denote the standardized predicted expression level of gene *g* across *N* individuals. Similarly, let *C_l_* ∈ R*N×*1 denote the observed value of the *l*th standardized covariate. We fit the following elastic net model.

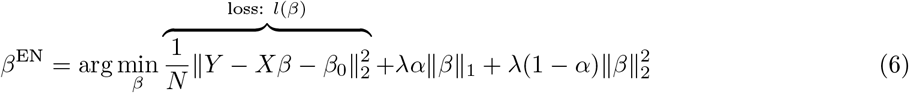

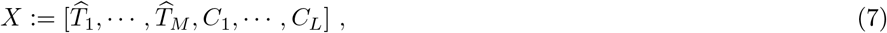

where *β*_0_ is the intercept, *M* is the number of genes, *L* is the number of covariates, 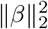 is the *l*_2_ norm and ∥*β*∥_1_ is the *l*_1_ norm of the effect size vector. Here, *α* controls the relative contribution of the *l*_1_ penalization term and λ controls the overall strength of regularization.

The model fitting procedure was implemented using tensorflow v2 with mini-batch proximal gradient method and the code is available at https://github.com/liangyy/ptrs-tf. We trained models at *α* = 0.1 (*α* = 0.5 and 0.9 show similar performance). And fixing the *α* value, as suggested in (Friedman et al., 2010){friedman:2010}, we trained a series of models for a sequence of λ’s starting from the highest. The maximum λ value, λ_max_, was determined as the smallest λ such that eq. (8) is satisfied.

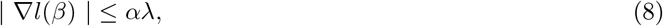

where the gradient is evaluated at

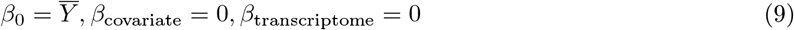

So, at λ = λ_max_, eq. (9) is the solution to eq. (6), which could serve as the initial points for the subsequent fittings of λ’s. We estimated λ_max_ using the first 1000 individuals of the data. And the sequence of λ was set to be 20 equally spaced points in log scale with the maximum value being 1.5λ_max_ and the minimum value being λ_max_*/*10^4^. Among these PTRS models generated at different λ values, we only kept the first 11 non-degenerate PTRS models so that we have the same number of candidate models for both PRS and PTRS.

### Transcriptome prediction models for PTRS construction

The predicted transcriptome in the discovery set (UKB EUR) was calculated using models from GTEx (Barbeira et al., 2020b) and MESA EUR based models (Mogil et al., 2018) listed in Supplementary Table S3). The GTEx EUR whole blood transcriptome consisted of 7,041 genes. For the MESA transcriptomes, we restricted the prediction to the 4,041 genes that were present in both the MESA EUR models and the MESA AFHI models (to ensure that PTRS built in the discovery set with the EUR models could be computed without missing genes in the target sets using the AFHI models).

### 1.9 Testing of PTRS in target sets

At the testing stage, given the standardized (within the population) predicted transcriptome of an individual, we calculated the PTRS of the individual using the following:

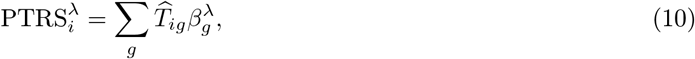

where *β*^λ^ denotes the *β*^EN^ obtained at hyperparamter equal to λ. For the PTRS built upon from GTEx EUR predicted transcriptome, the target PTRS was calculated with the GTEx EUR transcriptome (transcriptome predicted with GTEx EUR gene expression prediction models). To examine the utility of population-matched prediction model, the PTRS on the target set were calculated with of both MESA EUR and MESA AFHI transcriptomes.

### 1.10 Quantifying the prediction accuracy of PRS and PTRS with partial R2

To measure the predictive performance of PRS and PTRS, we calculated the partial *R*^2^ of PRS/PTRS against the observed phenotype accounting for the set of covariates listed in section 1.3. Specifically, for individual *i*, let 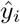 denote the predicted phenotype which could be either PRS or PTRS and *y_i_* denote the observed phenotype. Partial *R*^2^ (denoted as 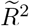 below) is defined as the relative difference in sum of squared error (SSE) between two linear models: 1) *y* ∼ 1 + covariates (null model); and 2) 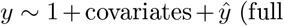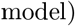 *i.e* 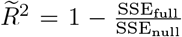. To enable fast computation, we calculated 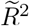 using an equivalent formula null shown in eq. (11) which relies on the projection matrix of the null model.

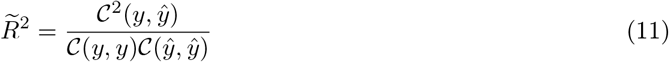

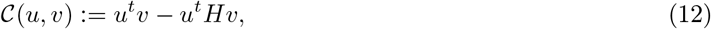

where *H* is the projection matrix of the null model, *i.e.* 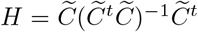 where 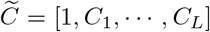 with *C_l_* being the *l*th covariate.

### 1.11 Quantifying the portability of PRS and PTRS

As stated in the results section, PTRS weights were computed in the discovery set (UKB EUR) and tested in the 5 target sets. For each of the 17 quantitative traits, 11 sets of weights for PRS and for PTRS were calculated with different hyperparameters. For PRS, different p-value thresholds were used to generate 11 different sets of weights. For PTRS, 11 different regularization parameters were used to generate the different sets of weights. The prediction accuracy in each of the 5 the target sets were calculated using the partial *R*^2^ described in section section 1.10 and the highest *R*^2^ among the 11 sets weights were used as the prediction accuracy. Portability was defined as the ratio of the prediction accuracy in each target set divided by the prediction accuracy in the European reference set. Therefore, by definition, portability in the EUR ref. set was 1.

When calculating the portability of PTRS using MESA AFHI transcriptome, we used the MESA EUR model 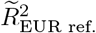 as the reference. This is a conservative choice since MESA EUR model is expected to perform better than MESA AFHI model among EUR individuals.

## Supporting information

Large Supplemental Tables

## Acknowledgements

We thank Owen Melia for help managing the UK Biobank data. This research has been conducted using the UK Biobank Resource under Application Number 19526. HKI and YL were partially funded by R01MH10766 and P30 DK20595 (Diabetes Research and Training Center).

## Disclosure

H.K.I. has received speaker honoraria from GSK and AbbVie.

## Code and data availability

Code for data analysis is at https://github.com/liangyy/ptrs-ukb. Code for mini-batch elastic net is at https://github.com/liangyy/ptrs-tf. The chip heritability and PVE analysis results are at Supplementary Table S4 and Supplementary Table S5. And the PRS and PTRS *R*^2^ results are at Supplementary Table S6 and Supplementary Table S7.

## Notes

### Summary of Updates

Acknowledgment of UK Biobank use

