## Supplemental Material for "Polygenic transcriptome risk scores improve portability of polygenic risk scores across ancestries"

November 9, 2020

### 1 Supplementary tables

| UKB Field Description | UKB Field ID | Tag | Phenotype Category |
|---|---|---|---|
| Standing height | 50 | Height | Height |
| Diastolic blood pressure, automated reading | 4079 | DBP | Blood pressures |
| Systolic blood pressure, automated reading | 4080 | SBP | Blood pressures |
| Body mass index (BMI) | 21001 | BMI | BMI |
| White blood cell (leukocyte) count | 30000 | WBC | Blood cell counts |
| Red blood cell (erythrocyte) count | 30010 | RBC | Blood cell counts |
| Haemoglobin concentration | 30020 | Hb | Haemoglobin related |
| Haematocrit percentage | 30030 | Ht | Haemoglobin related |
| Mean corpuscular volume | 30040 | MCV | Haemoglobin related |
| Mean corpuscular haemoglobin | 30050 | MCH | Haemoglobin related |
| Mean corpuscular haemoglobin concentration | 30060 | MCHC | Haemoglobin related |
| Platelet count | 30080 | Platelet | Blood cell counts |
| Lymphocyte count | 30120 | Lymphocyte | Blood cell counts |
| Monocyte count | 30130 | Monocyte | Blood cell counts |
| Neutrophill count | 30140 | Neutrophil | Blood cell counts |
| Eosinophill count | 30150 | Eosinophil | Blood cell counts |
| Basophill count | 30160 | Basophil | Blood cell counts |

Supplementary Table 1: **Meta information of the phenotypes retrieved from UK Biobank which were used in the analysis**. The "Tag" column shows the short name of the phenotyes used in this paper. And phenotypes are assigned into five categories which are shown in "Phenotype Category" column

| Ancestry | Number of individuals |
|---|---|
| AFR | 2835 |
| EUR | 356476 |
| E.ASN | 1326 |
| S.ASN | 4789 |

Supplementary Table 2: **Number of individuals included in the analysis stratified by ancestry**.

| Method | Data source | Population | Tissue | Number of genes | Sample size | Tag |
|---|---|---|---|---|---|---|
| CTIMP | GTEx V8 | European | Adipose_Subcutaneous | 9228 | 491 | |
| CTIMP | GTEx V8 | European | Artery_Tibial | 9027 | 489 | |
| CTIMP | GTEx V8 | European | Breast_Mammary_Tissue | 8127 | 337 | |
| CTIMP | GTEx V8 | European | Cells_Cultured_fibroblasts | 8731 | 417 | |
| CTIMP | GTEx V8 | European | Lung | 8954 | 444 | |
| CTIMP | GTEx V8 | European | Muscle_Skeletal | 7671 | 602 | |
| CTIMP | GTEx V8 | European | Nerve_Tibial | 10184 | 449 | |
| CTIMP | GTEx V8 | European | Skin_Sun_Exposed_Lower_leg | 9474 | 517 | |
| CTIMP | GTEx V8 | European | Thyroid | 9827 | 494 | |
| CTIMP | GTEx V8 | European | Whole_Blood | 7041 | 573 | GTEx EUR |
| Elastic Net | MESA | | Monocyte | 4670 | 578 | MESA EUR |
| Elastic Net | MESA | African American or Hispanic | Monocyte | 5554 | 585 | MESA AFHI |

Supplementary Table 3: **Meta information of the prediction models used in the analysis**. The highlighted prediction models were used to build PTRS. The "Tag" column shows the short name of the models used in this paper.

### 2 Supplementary tables in spreadsheet

The tables are available at
https://docs.google.com/spreadsheets/d/1wm9xVOa0hNtoEn5mS1fwlKtOuOvVOd7ODFiSBmFSJ9Q

Supplementary Table 4: **Chip heritability of the 17 quantitative traits in UK Biobank.**

Supplementary Table 5: **The PVE of predicted transcriptome of the 17 quantitative traits in UK Biobank.**

Supplementary Table 6: **Prediction performance of PRS for the 17 quantitative traits in UK Biobank.** The best $\widetilde{R}^2$ among all PRS candidates for each target set is listed.

Supplementary Table 7: **Prediction performance of PTRS for the 17 quantitative traits in UK Biobank.** The best $\widetilde{R}^2$ among all PTRS candidates for each target set is listed.
